# “Loss of alkyladenine DNA glycosylase alters gene expression in the developing mouse brain and leads to reduced anxiety and improved memory”

**DOI:** 10.1101/2023.10.05.561113

**Authors:** Diana L. Bordin, Kayla Grooms, Nicola P. Montaldo, Sarah L Fordyce Martin, Pål Sætrom, Leona D. Samson, Magnar Bjørås, Barbara van Loon

## Abstract

Neurodevelopment is a tightly coordinated process, during which the genome is exposed to spectra of endogenous agents at different stages of differentiation. Emerging evidence indicates that DNA damage is an important feature of developing brain, tightly linked to gene expression and neuronal activity. Some of the most frequent DNA damage includes changes to DNA bases, which are recognized by DNA glycosylases and repaired through base excision repair (BER) pathway. The only mammalian DNA glycosylase able to remove frequent alkylated DNA based is alkyladenine DNA glycosylase (Aag, aka Mpg). We recently demonstrated that, besides its role in DNA repair, AAG affects expression of neurodevelopmental genes in human cells. Aag was further proposed to act as reader of epigenetic marks, including 5-hydroxymethylcytosine (5hmC), in the mouse brain. Despite the potential Aag involvement in the key brain processes, the impact of Aag loss on developing brain remains unknown. Here, by using Aag knockout (*Aag*^*-/-*^) mice, we show that Aag absence leads to reduced DNA break levels, evident in lowered number of γH2AX foci in postnatal day 5 (P5) hippocampi. This is accompanied by changes in 5hmC signal intensity in different hippocampal regions. Transcriptome analysis of hippocampi and prefrontal cortex, at different developmental stages, indicates that lack of Aag alters gene expression, primarily of genes involved in regulation of response to stress. Across all developmental stages tested aldehyde dehydrogenase 2 (*Aldh2*) emerged as one of the most prominent genes deregulated in Aag-dependent manner. In line with the changes in hippocampal DNA damage levels and the gene expression, adult *Aag*^*-/-*^ mice exhibit altered behavior, evident in decreased anxiety levels determined in the Elevated Zero Maze and increased alternations in the Elevated T Maze tests. Taken together these results suggests that Aag has functions in modulation of genome dynamics during brain development, important for animal behavior.

**Highlights:**

- Aag loss results in reduced DNA damage signal in developing hippocampus;
- 5hmC signal intensity is perturbed in hippocampal regions of *Aag*^*-/-*^ mice;
- Gene expression is altered in *Aag*^*-/-*^ hippocampus and prefrontal cortex;
- Aag represses *Aldh2* expression;
- *Aag*^*-/-*^ mice have reduced anxiety and improved memory.

## INTRODUCTION

Throughout a lifetime, genomes are subjected to spectra of endogenous and exogenous agents that induce various types of DNA damage. Increase in DNA damage levels is generally thought to counteract genome stability and characterize common pathologies, aging and aging-associated processes like neurodegeneration [1]. Despite this, an increasing amount of evidence suggests that DNA damage and repair are not solely triggered by pathological processes [2-7]. The same DNA damage and repair processes involved in neurodegeneration, are proposed to have key roles in fundamental neurodevelopment and physiological neuronal functions, related to neural plasticity and activity [8, 9].

One of the most frequent types of DNA damage are DNA base lesions, arising from oxidation, deamination and alkylation [10]. Base excision repair (BER) oversees the removal of damaged and modified DNA bases, through the action of DNA glycosylases that act on specific substrates, thus determining pathway specificity. Alkyladenine DNA glycosylase (AAG, also known as MPG) is the only mammalian glycosylase able to remove frequent aberrantly methylated bases, including 7-methylguanine (7mG) and 3-methyladenine (3mA), as well as hypoxanthine (Hx) and 1,N^6^-ethenoadenine (εA), the byproducts of lipid peroxidation [11]. It is estimated that 7x10^3^ to 7x10^4^ aberrantly methylated DNA bases are generated in each differentiated cell, such as neuron, every day [12]. Substrate removal by Aag results in the generation of apurinic/apyramidic (AP) site, which is cleaved by AP endonuclease 1 (APE1), the subsequent gap filled by DNA polymerase (pol) β, and nick sealed by a XRCC1/DNA ligase III. The inability to coordinate BER steps results in accumulation of toxic BER intermediates like AP-sites and single-strand breaks (SSBs). Presence of SSBs in close proximity on opposite DNA strands may lead to formation of a double-strand breaks (DSBs) and subsequent signaling mediated through ATM-dependent H2AX phosphorylation (γH2AX) [13]. Imbalanced BER was shown to have harmful impacts on essential cellular processes, including replication, energy metabolism and cell viability [14-16]. In line with this, a body of evidence shows that *Aag* over-expression in mice results in higher toxicity upon exposure to alkylating drugs, while loss of Aag has protective functions in different tissues, such as the cerebellum and retina [17-21]. Besides the generation of toxic intermediates due to pathway imbalance, DNA glycosylases can be utilized in a programmed way [9, 22, 23]. In neurons, programed generation of SSBs, through activity of thymine DNA glycosylase (TDG), has been recently proposed to be required for physiological epigenetic (re)programming and activation of transcriptional programs allowing unperturbed neuronal differentiation [22]. In addition to TDG, other DNA glycosylases can impact gene expression through different mechanisms, many of which remain though only partially understood [24-26]. Using human cell lines, we showed that AAG associates with actively transcribing RNA pol II and modulates expression of neurodevelopmental genes [27]. In addition to complex formation with RNA pol II, AAG was demonstrated to interact with several transcription factors (TFs) like methylated DNA-binding domain 1 (MBD1), to potentially inhibit transcription [28]. AAG also associates with the estrogen receptor (ER) α. This interaction stimulates AAG glycosylase activity and stabilizes the binding of ERα to estrogen-response-elements (ERE) *in vitro* [29]. Independent of its glycosylase activity, Aag was suggested to act as reader of epigenetic DNA marks 5-hydroxymethylcytosine (5hmC) and 5-formylcytosine (5fC) [30, 31] in mouse stem cells and brain. 5hmC is especially abundant in the brain and embryonic stem cells, and impaired 5hmC levels in the mouse brain were reported to interfere with spatial learning and memory [32]. Despite Aag impacts on neurodevelopmental gene expression and ascribed functions as reader of epigenetic DNA marks, *in vivo* roles of Aag in the brain development and behavior remain unknown.

In this study, by analyzing different brain regions, isolated from wild-type (WT) and Aag knock-out (*Aag*^*-/-*^) mice, we demonstrate the impact of Aag on DNA damage and 5hmC levels. Specifically, loss of Aag led to reduction in the number of γH2AX foci across hippocampus at postnatal day 5 (P5), accompanied by changed 5hmC signal in various hippocampal regions. To further elucidate the involvement of Aag in gene expression during neurodevelopment and its possible consequences in adult life, we performed RNA sequencing analysis in WT and *Aag*^*-/-*^ hippocampi and prefrontal cortex at P5, 6 weeks (6W) and 6 months (6M). Importantly, loss of Aag altered gene expression in both hippocampus and prefrontal cortex, across all analyzed developmental stages, with most prominent effects being observed for genes involved in regulation of response to stress at P5. Across all developmental stages tested, aldehyde dehydrogenase 2 (*Aldh2*) was identified as one of the most significant Aag-dependent genes. The changes in hippocampal gene expression, 5hmC and DNA damage status were accompanied by altered behavior in adult *Aag*^*-/-*^ mice, evident in decreased anxiety and increased alternations in Elevated Zero Maze and Elevated T Maze tests, respectively. In summary, our findings represent the first evidence of Aag impacts on genome dynamics and gene expression in developing brain *in vivo*, relevant for brain functioning and consequently animal behavior.

## RESULTS AND DISCUSSION

### Loss of Aag results in reduced number of γH2AX foci and altered 5hmC signal intensity in the hippocampal regions of P5 mice

Generation of DNA breaks through BER activity has emerged as important determinant of neuronal development, activity and synaptic plasticity [6, 7, 33]. To address the impact of Aag loss on status of DNA breaks in the developing mouse brain, immunofluorescence (IF) analysis of γH2AX foci was performed in P5 WT and *Aag*^*-/-*^ hippocampal regions (Fig. 1A). We focused on the hippocampus, a site of continuous neurogenesis and with central roles in the storage of memories, spatial memory and emotional responses [34]. On the level of whole hippocampus, the number of endogenous γH2AX foci was significantly lower in *Aag*^*-/-*^ nuclei, when compared to WT (Fig. 1B). Similarly, the number of endogenous γH2AX foci was previously shown to be reduced in *Aag*^*-/-*^ mouse liver and fibroblasts [35, 36]. To determine whether the observed effect was global, or specific to defined hippocampal regions, number of γH2AX foci was analyzed next in the dentate gyrus (DG), cornum ammonis (CA) 1 and CA3 regions. DG is the neurogenic region of the hippocampus, rich in granular neurons, while CA1 and CA3 regions are formed by a mix of excitatory and inhibitory pyramidal neurons. Interestingly, the number of γH2AX foci was significantly reduced in *Aag*^*-/-*^ DG and CA1, while no significant change was observed in the CA3 region (Fig. 1C-E), thus suggesting region-specific effects of Aag loss. The status of γH2AX was shown to positively associate with proliferation and neurogenesis capacity in embryonic, postnatal and adult mouse brain, as well as in neural stem cells [37, 38]. To test if the detected γH2AX differences in *Aag*^*-/-*^ hippocampi resulted from altered proliferation, Ki67 status was analyzed. No significant difference in the Ki67 signal intensity were detected between WT and *Aag*^*-/-*^ DG (Fig. S1A-B). This suggests that the level of γH2AX foci directly relates to Aag status in DG at P5.

**Figure 1.**
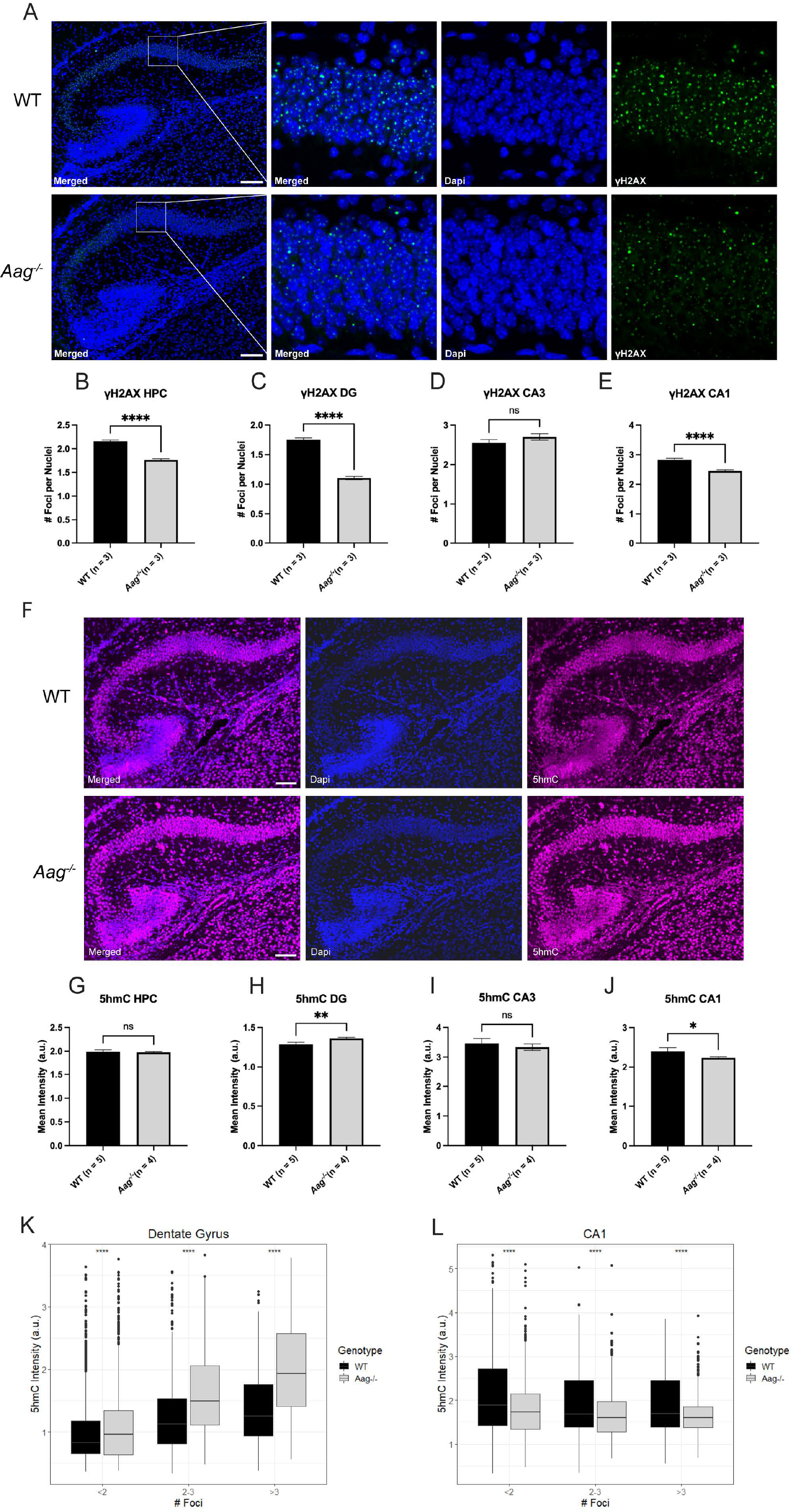
Number of γH2AX foci is reduced and 5hmC signal intensity changed in different hippocampal regions of *Aag*^*-/-*^ mice, at age P5. (A) Representative immunofluorescence images of the γH2AX signal in the hippocampus (HPC) of WT and *Aag*^*-/-*^ mice at postnatal day 5 (P5) (Scale bars = 100 μm). (B-E) Quantification of number of γH2AX foci per cell nucleus in (B) total HPC and in the specific HPC regions: (C) dentate gyrus (DG), (D) CA3 and (E) CA1. 1-2 hemispheres from 3 animals were included; data represented as mean ± SEM; statistical analysis performed using two-tailed Student’s t-test: * p < 0.05, **p < 0.01, **** p < 0.0001 in (B-E). (F) Representative 5hmC staining of P5 WT and *Aag*^*-/-*^ mice HPC. (G-J) Quantification of 5hmC intensity per cell normalized to the Dapi intensity of the same cell within: (G) total HPC and the HPC regions (H) DG, (I) CA3, and (J) CA1, respectively. 1-2 hemispheres from 4-5 animals were analyzed; data represented as mean ± SEM; statistical analysis performed using one-tailed Student’s t-test: * p < 0.05, **p < 0.01, **** p < 0.0001 in (G-J). (K and L) Analysis of signal intensity of 5hmC binned by the number of γH2AX foci per cell in the DG (K) and CA1 (L) regions of n = 3 hippocampi of WT and *Aag*^*-/-*^ mice at P5. Data represented as median and quartiles, statistics performed using two-tailed Student’s t-test: **** p < 0.0001.

In addition to its role in DNA repair, Aag was proposed to act as a reader of epigenetic DNA modifications, like 5hmC, with critical functions in neurodevelopment [39]. To test if lack of Aag affects 5hmC levels, we next performed IF analysis targeting 5hmC in WT and *Aag*^*-/-*^ hippocampi (Fig. 1F). While total 5hmC levels were unchanged in WT compared to *Aag*^*-/-*^ hippocampi, we detected a significant increase in 5hmC signal in DG and a decrease in CA1 region (Fig 1G-J). As both 5hmC and γH2AX status were altered in *Aag*^*-/-*^ DG and CA1, and since 5hmC was previously shown to accumulate at the sites of γH2AX foci [40], we next assessed the relation between 5hmC signal intensity and the number of γH2AX foci per cell nucleus (Fig. 1K-L). Interestingly, a linear relationship between the number of γH2AX foci and 5hmC signal intensity was prominent and specifically observed in the DG, but not CA1 and CA3 regions (Fig. 1K-L and Fig. S1C). As expected 5hmC signal intensity was higher in *Aag*^*-/-*^ DG nuclei and lower in CA1 nuclei; this effect was irrespective of the number of γH2AX foci (Fig. 1K-L). The observed changes could have functional impacts, since induction of γH2AX foci was shown to associate with induction of early-response genes and to accompany responses to normal behavior in defined brain regions, primarily DG [2, 3]. Perturbed 5hmC status and associated changes in the gene expression programs were similarly demonstrated to affect memory and learning in mice [41]. Taken together, the detected changes in γH2AX and 5hmC status suggest possible differences in *Aag*^*-/-*^ gene expression and brain function.

### Gene expression is deregulated in *Aag*^*-/-*^ hippocampus and prefrontal cortex across developmental timepoints

We previously identified AAG as an important modulator of neurodevelopmental gene expression in human cells [27]. In line with this and the observed changes in γH2AX and 5hmC status in *Aag*^*-/-*^ hippocampal regions (Fig. 1), we tested the impact of Aag loss on gene expression *in vivo* at different developmental stages. RNA-sequencing analysis identified sets of differentially expressed genes (DEGs) in both *Aag*^*-/-*^ hippocampal and prefrontal cortex samples isolated at P5, 6W and 6M (Fig. 2A-C and S2A-C), thus indicating a potential role for Aag in gene expression regulation across multiple time points. Interestingly, most DEGs in *Aag*^*-/-*^ hippocampi were observed at P5, suggesting that the Aag-mediated effect on gene expression programs is broadest during early development. Subsequent comparison of P5 *Aag*^*-/-*^ and WT hippocampal transcriptomes indicated that majority of 240 DEGs belonged to the cellular stress response process (Fig. 2D). Interestingly, the same biological process was also identified as the most significant in the *Aag*^*-/-*^ prefrontal cortex DEGs (Fig. S2D). These results, together with the recent work that identified Aag as a merging point for different stress response processes [42], propose roles of Aag in cellular stress responses, both in hippocampus and prefrontal cortex at P5. To further determine genes deregulated by Aag loss at all developmental time points, we next compared the P5, 6W and 6M transcriptomes. The heat maps presented in Fig. 2E and S2E reveal four commonly DEGs during the three developmental periods, both in hippocampus and prefrontal cortex, thus suggesting significant Aag involvement in the regulation of those genes. The expression of *Aldh2* (aldehyde dehydrogenase 2) and *Gabra2* (γ-aminobutyric acid receptor subunit α-2) was upregulated, while *Pttg1* (PTTG1 regulator of sister chromatid separation, securin) and *Ublcp1* (ubiquitin like domain containing CTD phosphatase 1) were downregulated, in *Aag*^*-/-*^ hippocampus and prefrontal cortex across the time points (Fig. 2E and S2E). The subsequent qPCR analysis of Aldh2, Gabra2 and Pttg1 mRNA levels in *Aag*^*-/-*^ hippocampi, confirmed the RNA-seq results (Fig. 2F-H). Ublcp1 mRNA levels were however not significantly different between WT and *Aag*^*-/-*^ samples (Fig. S2F and G). The observed deregulation of *Aldh2* and *Gabra2* expression points at possibly wider BER contribution, since lack of XRCC1 was similarly shown to result in ALDH2 upregulation in human cells [43]. Further, loss of murine DNA glycosylases Neil 1, 2 and 3 altered *Gabra2* expression, however in contrast to Aag, *Gabra2* was downregulated in Neil deficient brains [44, 45].

**Figure 2.**
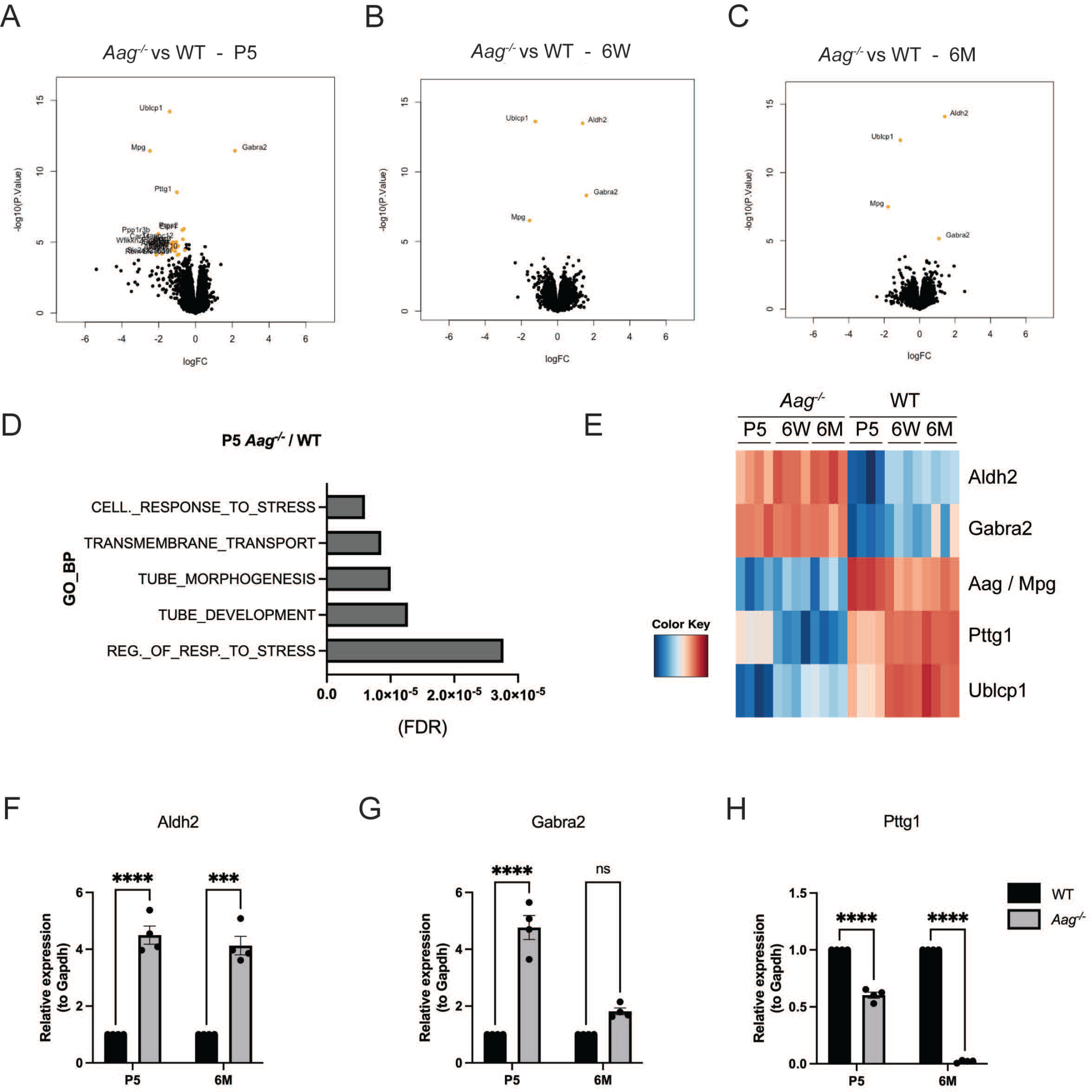
Loss of Aag alters gene expression in mice hippocampi across different developmental stages. (A-C) Volcano plots depicting deferentially expressed genes (DEGs) in *Aag*^*-/-*^ compared to WT hippocampus, at: (A) postnatal day 5 (P5), (B) 6 weeks (6W) and (C) 6 months (6M). Orange dots represent specific significant DEGs with ≥1.5-fold change and FDR<0.05. (D) Top five biological processes (BP) gene ontology (GO) terms determined by the Gene Set Enrichment Analysis (GSEA) for all DEGs, at an FDR<0.2, in *Aag*^*-/-*^ compared to WT P5 hippocampus. (E) Heat map of the expression levels of genes differentially expressed in the *Aag*^*-/-*^ compared to WT hippocampus at all tested ages (P5, 6W, and 6M). Color scale is representing change in the gene expression depicting log2 fold change relative to the mean. (F-H) RT-qPCR quantification of (F) Aldh2, (G) Gabra2 and (H) Pttg1 mRNA levels in the hippocampi of WT and *Aag*^*-/-*^ mice at P5 and 6M (n=4). Statistics performed using two-way ANOVA with Šídák′s multiple comparison test: *** p < 0.001; **** p < 0.0001.

### Expression of *Aldh2* is directly impacted by Aag in P5 hippocampi

To elucidate to which extent Aag directly modulates *Aldh2, Gabra2* and *Pttg1* expression, we analyzed the mRNA levels in P5 WT, *Aag*^*-/-*^, and Aag overexpressing (*Aag*^*Tg*^) hippocampal samples. While Aldh2 was as expected upregulated in *Aag*^*-/-*^, the hippocampi overexpressing Aag exhibited significant *Aldh2* downregulation compared to WT (Fig 3A). In contrast to *Aldh2*, the expression of *Pttg1* and *Gabra2* was not directly dependent on Aag status (Fig. S3). The expression of *Pttg1* was unchanged in *Aag*^*-/-*^ and *Aag*^*Tg*^, while *Gabra2* expression was comparable in *Aag*^*Tg*^ and WT hippocampi. Taken together, these results suggests that Aag specifically regulates *Aldh2*, but not *Gabra2* and *Pttg1*.

**Figure 3.**
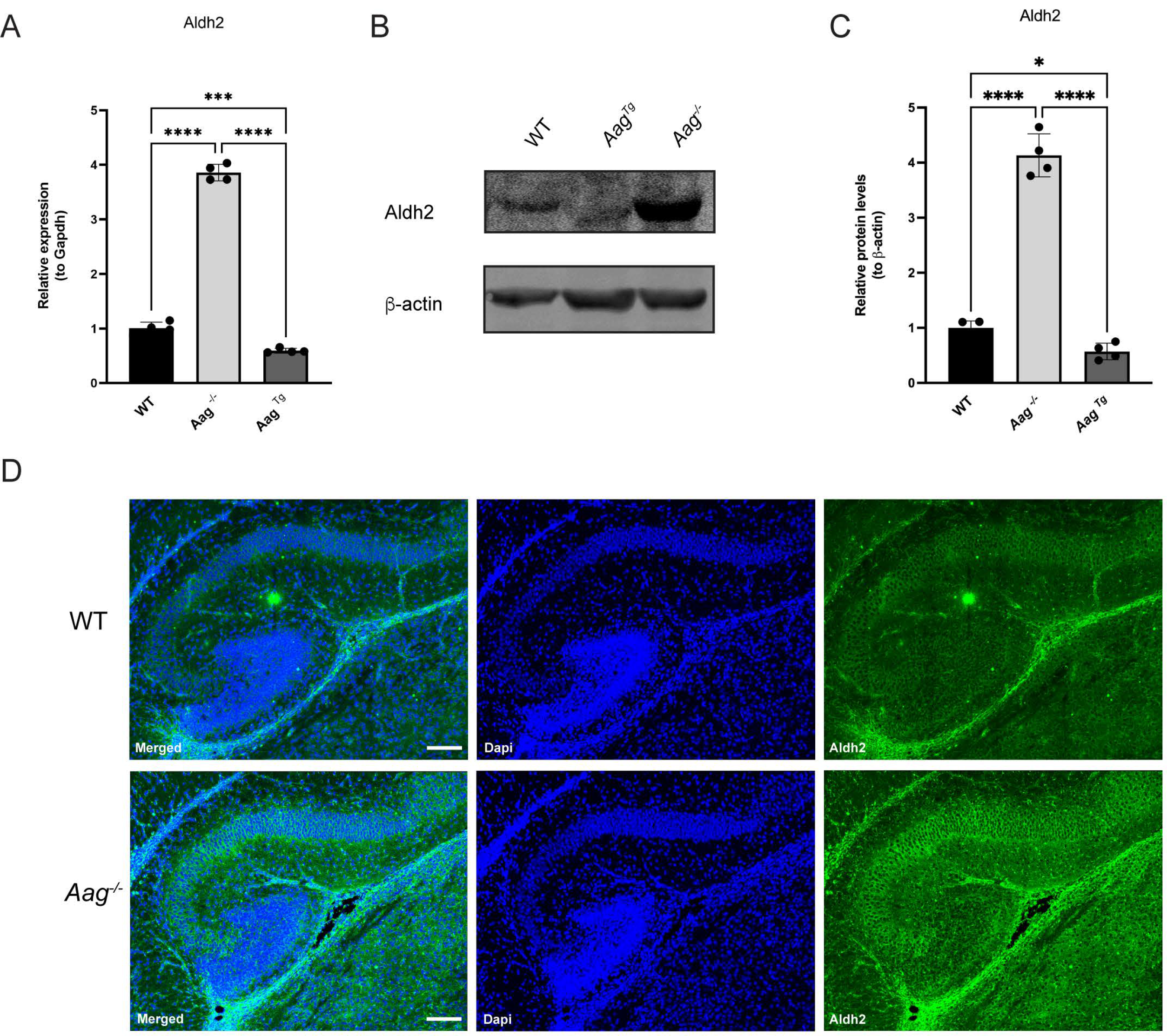
Aldh2 mRNA and protein levels are regulated in Aag-dependent manner in P5 hippocampi. (A) RT-qPCR analysis of Aldh2 mRNA levels in the hippocampus of WT, *Aag*^*Tg*^, and *Aag*^*-/-*^ at postnatal day 5 (P5) (n=4). (B) Western blot analysis of Aldh2 levels in P5 WT, *Aag*^*Tg*^, and *Aag*^*-/-*^ mouse hippocampi. (C) Quantification of four independent experiments as the one depicted in (B). (D) Representative immunofluorescence images of Aldh2 staining within the hippocampus of WT and *Aag*^*-/-*^ mice at P5 (Scale bars = 100 μm). Data represented as mean ± SEM, statistics performed using one-way ANOVA with Holm-Šídák′s multiple comparison test: * p < 0.05, *** p < 0.001, **** p < 0.0001.

Aldh2 is the key enzyme in the oxidative metabolism of toxic aldehydes in the brain (as 4-hydroxynonenal (4-HNE), product of lipid peroxidation) and has important functions in sanitizing endogenous formaldehyde, a genotoxic metabolite of many cellular processes [46-48]. Due to this, and the high Aldh2 presence in different brain regions [49], we tested to which extent Aag-mediated *Aldh2* regulation causes relevant changes at the enzyme level. Immunoblot analysis targeting Aldh2 was performed in WT, *Aag*^*-/-*^ and *Aag*^*Tg*^ lysates from P5 hippocampi (Fig. 3B-C). We show that not only was *Adlh2* significantly overexpressed, but also Aldh2 protein levels were increased in the hippocampi of *Aag*^*-/-*^ mice. Accordingly, *Aag*^*Tg*^ hippocampi were characterized by reduced Aldh2 protein levels. The immunoblot results were further supported by immunofluorescence analysis revealing stronger Aldh2 signal in *Aag*^*-/-*^ when compared to WT P5 hippocampi (Fig. 3D). These findings thus indicate that Aag status impacts both Aldh2 mRNA and protein levels. The significant Aldh2 accumulation could have important neuroprotective impacts in *Aag*^*-/-*^ mice, since prior research showed that Aldh2 overexpression can reduced cell death in different models of frequent neurodegenerative diseases, such as Alzheimer′s and Parkinson′s disease [48]. Further in gastric mucosa cells ALDH2 was identified to be part of protective mechanism against endogenously produced 4-HNE from lipid peroxidation, and *ALDH2* overexpression in these cells was associated with a reduction in γH2AX levels [50]. Additionally, Aldh2 is known to have important impacts on cognition [48], thus the *Aldh2* overexpression could likely impact brain functions in *Aag*^*-/-*^ mice.

### Loss of Aag effects learning and memory in adult mice

The here observed changes in γH2AX and 5hmC status, altered gene expression profiles and specifically *Aldh2* overexpression in *Aag*^*-/-*^ hippocampus, collectively have potential to affect the brain performance. Given the importance of the hippocampus in cognitive functions, and its relevant roles in the emotional behavior, including the regulation of anxiety responses [34], we next assessed whether loss of Aag has functional implications and affects behavior in mice. To address this, WT and *Aag*^*-/-*^ mice at 4 to 6 months were first tested for anxiety levels and locomotor activity using elevated zero maze (EZM). Anxiety was evaluated considering the time spent in open arms. *Aag*^*-/-*^ mice spent significant more time exploring the open areas indicating decreased anxiety compared to WT animals (Fig. 4A). Additional locomotion measurements, assessed as the total distance travelled, indicated that *Aag*^*-/-*^ mice traveled significantly shorter distances, thus suggesting reduced activity (Fig. 4B). This could be either due to *Aag*^*-/-*^ mice spending more time pausing and exploring open arms (data not shown), or due to differences in locomotion, however this is not likely as no gross locomotory deficiencies had been detected in these mice [51]. Next, *Aag*^*-/-*^ mice underwent cognition and spatial working memory testing using the spontaneous alternation version of the T-maze. Mice lacking Aag showed significantly improved capacity of memorizing the visited arms in comparison with WT mice (Fig. 4C). Taken together, these results suggest that Aag is involved in modulation of anxiety-like behavior and memory. The observed reduced anxiety of *Aag*^*-/-*^ mice, could in part be caused by increased Aldh2 levels, since *Aldh2* overexpression was similarly shown to reduce anxiety and depressive-like behavior in rat models of depression [52]. *Aldh2* overexpression was further demonstrated to have beneficial impact on memory both during aging and in different neurodegenerative disease models [48].

**Figure 4.**
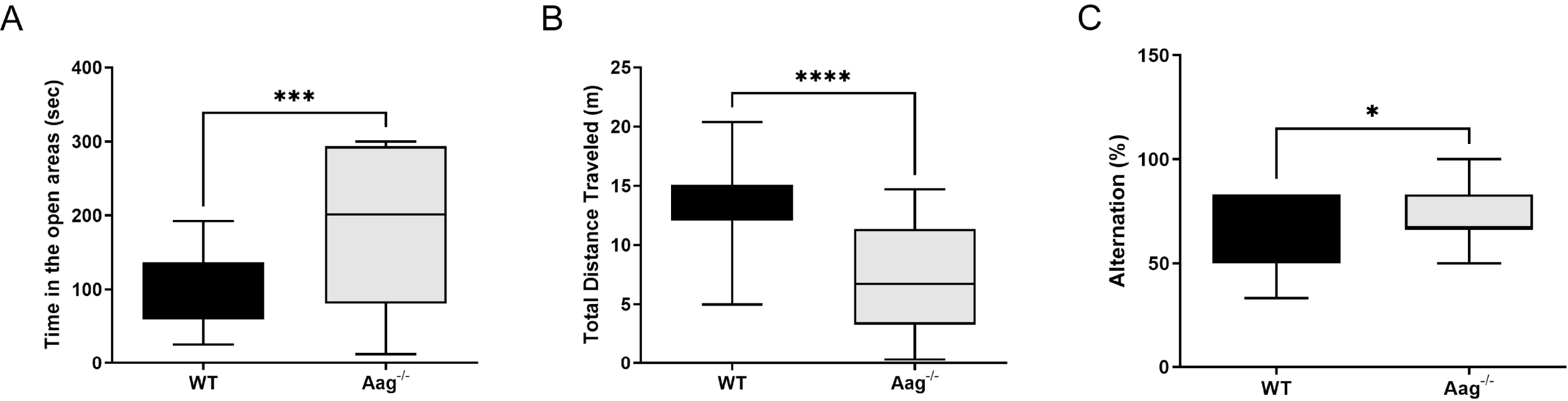
*Aag*^*-/-*^ mice exhibit reduced anxiety and improved memory. (A-C) Elevated zero maze (EZM) and T-maze testing of adult (4-6 month old) WT (n= 29) and *Aag*^*-/-*^ (n= 23) mice behavior. (A) Time spent (seconds) in open arms of the EZM. (B) Total distance traveled (meters) in EZM. (C) Percentage of spontaneous alternation in T-maze test. Statistical analysis was performed using two-tailed Student’s t-test: * p < 0.05; *** p < 0.001, **** p < 0.0001.

In summary, our results suggest that loss of Aag, has a multifaced impact, affecting hippocampal genome integrity, epigenome and gene expression profiles, relevant for cognition and behavior. While future mechanistic insights are needed, the here identified Aag-Aldh2 axis emerged as one of the central points in these processes, with potential to both contribute to reduction in γH2AX foci, as well as reduced anxiety and improved memory phenotypes.

## MATERIALS AND METHODS

### Mice

C57BL/6J, used as WT, *Aag*^*-/-*^ and *Aag*^*Tg*^ mice were generated as described in (Engelward et al., 1997). To characterize brain development in *Aag*^*-/-*^ versus WT, mice at postnatal day five (P5), juvenile mice at six weeks (6W) and adult mice from four to six months (6M) were evaluated. All animals were maintained in pathogen-free conditions at the Comparative Medicine Core Facility of the Norwegian Science and Technology and the experiments were approved and conducted in accordance with the Norwegian Animal Research Authority.

### Mouse genotyping

WT, *Aag*^*-/-*^ and *Aag*^*Tg*^ were genotyped according to [51]. Sex determination of pups at postnatal day 5 was made by the amplification of the Y chromosome gene *Sry*. PCR products from mice were purified with QIAquick PCR Purification Kit (Qiagen, 28104) and the genotype was confirmed by DNA sequencing. All primer sequences are listed in Supplementary table 1.

### Hippocampus and pre-frontal cortex isolation

Hippocampus and pre-frontal cortex were isolated from P5, 6W and 6M male mice (4 animals per time point) for further RNA extraction. Brains were immediately removed after the dislocation of the neck. The isolation of the brain regions was performed on ice and stored in liquid nitrogen until the analysis.

### Tissue homogenization and total RNA isolation from different brain regions

Total RNA from hippocampus and pre-frontal cortex was purified with AllPrep DNA/RNA/Protein Mini Kit (QIAGEN, 80004) and DNAseI digested (QIAGEN, 79245). Frozen hippocampus and pre-frontal cortex were transferred to a 2 ml lysing tube (Precellys, KT03961-1-405.2) pre-filled with 1.4 mm zirconium oxide beads (Precellys, 03961-1-103) and 600 μl of RLT buffer (provided by the AllPrep DNA/RNA/Protein Mini Kit). Samples were then homogenized with MagNAlyser (Roche Diagnostics GmbH) at 5000 rpm during 15 sec and finally centrifuged at full speed for 3 min. Supernatant was transferred to an AllPrep spin column and isolation was performed according to the manufacturer’s protocol.

### Library preparation and RNA sequencing

RNA libraries were built using the Lexogen SENSE mRNA library preparation kit (polyA stranded). Libraries were sequenced using stranded pair-end 75bp sequencing on the Illumina HiSeq 4000, with each sample being sequenced across all 8 lanes. Illumina bcl2fastq 2.20.0.422 was used for base calling. Sequenced reads were mapped to GRCm38.6 whole genome using STAR v2.4.0 with parameters --chimSegmentMin 30 --runThreadN 12 –outFilterMultimapNmax 20 --alignSJoverhangMin 8 --alignSJDBoverhangMin 1 --outFilterMismatchNmax 10 --outFilterMismatchNoverLmax 0.04 --alignIntronMin 20 --alignIntronMax 1000000. Counting was performed using HTSeq [53] v0.6.0 (htseq-count) using reverse stranded settings, counting exon features and reporting Ensembl Gene Ids. Data normalization and differential expression analysis was performed using limma v3.32.10 [54], in R v 3.4.1, filtering out genes with expression (total normalized read counts) less than the number of samples.

### Gene expression analysis

Total RNA from tissue samples was reverse transcribed using High-Capacity cDNA Reverse Transcription Kit (Applied Biosystems, 4368813) according to the manufacturer’s protocol. qPCR was performed with Power SYBR Green PCR Master Mix (ThermoFisher Scientific, 4368708) on a StepOnePlus Real-Time PCR System (ThermoFisher Scientific, 4376600). Relative transcription levels were determined by normalizing to GAPDH mRNA levels. All primer sequences are listed in Supplementary table 1.

### Immunoblot analysis

Total proteins from P5 hippocampi were purified with AllPrep DNA/RNA/Protein Mini Kit (Qiagen, 80004). The protein pellets were then resuspended in 90 μl Buffer ALO (Qiagen) supplemented with 0.4X NuPAGE™ LDS Sample Buffer (Thermofisher, NP0007). Samples were sonicated using Bioruptor® Pico (Diagenode, B01060010) for 5 minutes (30-second ON and 30-second OFF cycle). The samples were then incubated for 15 minutes at 95 °C, followed by 5 minutes at RT. Residual insoluble material was pelleted by centrifugation at 14,000 g for 2 minutes. For SDS-PAGE, 20 μl of the protein samples were loaded onto NuPage 4–12% Bis–Tris polyacrylamide gels (Invitrogen, NP0321) and separated via electrophoresis. Subsequently, the proteins were transferred to 0.45 μm PVDF membranes in transfer buffer (192 mM glycine, 25 mM Tris base, 5% methanol). The primary antibodies used for immunoblotting analysis were anti-Aldh2 (Invitrogen, MA5-17029, 1:1000) and anti-β-actin (Sigma, A1978, 1:10000), followed by detection using infrared (IR) Dye-conjugated secondary antibodies (Li-COR Biosciences, 926-32210, 1:15000). The signal was captured using an Odyssey Scanner (LI-COR Biosciences). Protein levels were quantified using ImageJ software.

### Immunofluorescence

Brains from P5 mice were obtained after decapitation and immediately immersion fixed in 4% PFA. Sagittal brain sections of 9 μm of thickness were prepared using the microtome LeicaRM2255 (Leica Biosystems) and stored at 4 °C until use. Before the immunostaining, the sections were deparaffinized through incubation at 56 °C for 15 min and washed for 5 min x2 with Clear-Rite. Next, the sections were hydrated for 3 min x2 in 100% ethanol, 2 min in 96% ethanol, 1 min in 70% ethanol and water. Heat-induced antigen retrieval was performed in citrate buffer (0.01 M, pH 6.0) using a pressure cooker for 3 min. Slides were then cooled down at RT, permeabilized with 0.25%Triton X-100/PBS for 15 min and blocked for 1 h at RT with 5% NGS (Jackson Immuno Research, 005-000-121) + 5% BSA (Sigma, A2153). The following primary antibodies were used: Aldh2 (1:200 ThermoFisher Scientific, MA5-17029); 5hmC (1:250, Active Motif, 39769); γH2AX (Ser 139) (1:50, Merck 05-636); Ki67 (1:200, Abcam, ab15580). All primary antibody dilutions contained 0.5% NGS + 0.5% BSA for nonspecific reactivity. After the incubation overnight at 4 °C with primary antibodies, the brain sections were washed for 5 min x3 in 0.1% Tween 20/PBS and incubated in fluorescence tagged secondary antibodies for 2 hours at RT: Alexa Fluor 594 anti-rabbit (1:500, ThermoFisher Scientific, A11037), Alexa Fluor 488 anti-mouse IgG1 (1:500, Thermo Fisher Scientific, A21121), Alexa Fluor 647 anti-chicken (1:500, Abcam, ab150175. The brain sections were washed again for 5 min x3, counterstained with DAPI dilution (1:1000 in PBS, ThermoFisher Scientific, 62248) for 10 min and mounted with ProLong™ Gold Antifade Mountant with DAPI (ThermoFisher Scientific, P36931). Upon staining the sections were viewed using Zeiss 880 AiryScan using LD LCI Plan-Achromat 40 x/1.2 lmm AutoCorr DCI M27 objective. Images were acquired using Zen Black software (2.3 SP1). Tiling scan function with 10% overlap and Z-stack with a step size between 0.8 and 2μm was used to obtain 8 bit-rate images of the whole hippocampal region. Gain and step size were chosen for optimal signal to noise ratio and maintained within experiments.

### Image processing and analysis

Raw LSM5 files were opened and stitched in Fiji ImageJ (v.1.53f51, National Institutes of Health) with the Grid/Collection Stitching plug-in [55] using type: positions from file and order defined by image metadata. The imported 3D image was then flattened into a 2D representative image using the summed intensity Z projection. The entire hippocampus, dentate gyrus granule cell layer, CA3, and CA1 pyramidal cell layer regions were traced using the freehand tool to create regions of interest. Raw integrated density measurements of Ki67 within the region of interest were measured and normalized to the raw integrated density of the Dapi channel. 5hmC and γH2AX were analyzed using the StarDist 2D plugin for Fiji [56], using the pretrained model “Versatile (Fluorescent Nuclei)”. Probability and overlap thresholds were set for optimal cell counting with a Wildtype control image and all settings were maintained within experiments. Cells that were detected outside of the region of interest were deleted, and the raw integrated density of 5hmC signal within each cell was measured and normalized to the Dapi raw integrated density within the same cell. To count the number of γH2AX foci in the image, the Find Maxima tool was used with a single points output, and a Prominence value was maintained for all images within experiments. The raw integrated density of the points output was measured using the ROI output from StarDist and each value per cell was divided by 255 to obtain the number of points within each cell. Analysis of co-stained images with γH2AX and 5hmC was conducted in R (R Core Team (2021), v.4.1.3). Measurements for individual cells was imported, and outliers with a Z-score > 3 were removed. Cells were binned based on the number of foci and 5hmC raw integrated density normalized to Dapi for each cell was plotted with ggplot2 [57].

### Behavioral studies

Male mice four to six months old WT (n= 29) and *Aag*^*-/-*^ (n= 23) were used in all behavioral experiments presented herein. The researchers remained hidden behind a curtain during testing. Mice performing tasks were recorded and tracked at 10 Hz by the ANY-maze video tracking system (Stoelting, IL, USA).

### Elevated zero maze

The test was conducted in a custom-made apparatus (described in [58]) consisting of a black 5-cm-wide circular runway with four alternating open and closed areas. Animals were placed on the maze facing a closed arm and allowed to explore for 5 min. Anxiety levels were evaluated according to the time spent in open arms. The total distance travelled was used to measure the activity of mice.

### T-maze

Spontaneous alternation was evaluated in an enclosed T-maze consisting of three arms of equal size (30 x 10 x 10 cm) (Deacon and Rawlins, 2006). For each round of the experiment, the animals were placed in the starting arm of the apparatus and allowed to explore and choose one of the goal arms for ninety seconds. As soon as the animal entered the chosen arm, the door of the compartment was closed, and the mouse was confined for thirty seconds. After this period, the animal was gently removed and placed in a cage for one minute (retention time) before to start the second-choice trial. The test comprised in total 6 rounds performed during three consecutive days.

## Data availability

The RNA sequencing data reported in this paper are available in GEO under accession GSE243469 (https://www.ncbi.nlm.nih.gov/geo/query/acc.cgi?acc=GSE243469).

## Statistical Analysis

Statistical analysis was performed using GraphPad Prism (GraphPad Software, Inc., La Jolla, CA, v9.1.2) or R (R Core Team (2021), v.4.1.3). Significance was determined by two-tailed Student’s t-test in Fig. 1G-L, S1C and Fig. 4., one-tailed Student’s t-test in Fig. 1 B-E and S1B; two-way ANOVA with Šídák′s multiple comparison test in Fig. 2F-H and S2F-G and one-way ANOVA with Holm-Šídák′s multiple comparison test in Fig. 3A, C and S3.

## CRediT AUTHORSHIP DECLARATION STATEMENT

Diana L. Bordin: Investigation, Validation, Writing – original draft, Writing – review, Supervision. Kayla Grooms: Investigation, Validation, Writing – original draft, Writing – review. Nicola P. Montaldo: Investigation, Validation, Writing – editing. Sarah L Fordyce Martin: Validation, Writing – editing. Pål Sætrom: Validation, Supervision, Writing – editing. Leona D. Samson – Supervision, Writing – editing. Magnar Bjårøs – Conceptualization, Writing – editing. Barbara van Loon: Conceptualization, Supervision, Writing – original draft, Writing – review & editing.

## ACKNOWLEDGMENTS

This work has been supported by Onsager Fellowship and the Research Council of Norway grant (ID: 335324) to B.v.L. The RNA library preparation and RNA sequencing were performed at the Genomics Core Facility (GCF), while sectioning and microscopy at the Cellular and Molecular Imaging Core Facility (CMIC), Norwegian University of Science and Technology (NTNU). GCF and CMIC are funded by the Faculty of Medicine and Health Sciences (NTNU) and Central Norway Regional Health Authority. We thank to Rajikala Suganthan for training in hippocampal isolations; to Katja Scheffler, Nicole Bethge and Andreas Abentung for inputs and help with behavioral studies; to Hilde L. Nilsen for discussions and feedback to the manuscript; to current and past van Loon group members for discussions and suggestions.

