## Supplemental Info for "“Loss of alkyladenine DNA glycosylase alters gene expression in the developing mouse brain and leads to reduced anxiety and improved memory”"

**SUPPLEMENTARY TABLES**

**Table S1. List of DNA primers used to genotype mice and for RT-qPCR analysis of mRNA levels.**

| **Genotyping primers** | | |
|---|---|---|
| **Name** | **Forward / Reverse** | **Sequence 5' to 3'** |
| *Aag_PI (WT)* | Fw | GCAGCGACTGGCAGATTC |
| *Aag_PII (NEO) (Aag$^{-/-}$)* | Fw | AGAAGGGTGAGAACAGAG |
| *Aag_PIII (WT/Aag$^{-/-}$)* | Rv | GAAATGCACGGGCTAGGG |
| *Sry* | Fw | TTGTCTAGAGAGCATGGAGGGCCATGTCAA |
| | Rv | CCACTCCTCTGTGACACTTTAGCCCTCCGA |
| **cDNA primers** | | |
| **Name** | **Forward / Reverse** | **Sequence 5' to 3'** |
| *Gapdh* | Fw | GTCCCGTAGACAAAATGG |
| | Rv | CGCCCAATACGGCCAAA |
| *Aldh2* | Fw | TTCCCACCGTCAACCCTTC |
| | Rv | CCAATCGGTACAACAGCCG |
| *Gabra2* | Fw | AGAATCGGTGCCAGCAAGAA |
| | Rv | TTCGGGAGGGAATTTCGAGC |
| *Pttg1* | Fw | CTCCAACCAAAACAGCCGAC |
| | Rv | GGTAGGCATCATCAGGAGCA |
| *Ublcp1* | Fw | ACGCCAGAAGTTACTAGGGC |
| | Rv | CAAGCTCTCCTCTCGAGTTCC |

#### SUPPLEMENTARY FIGURE LEGENDS

**Figure S1. Impact of Aag loss on proliferation in the dentate gyrus, and the relation between 5hmC signal intensity and the number of γH2AX foci in the CA3 region of hippocampus.** (A) Representative immunofluorescence images of $Ki67^+$ cells in the dentate gyrus (DG) of postnatal day 5 (P5) WT and *Aag*$^{-/-}$ mice (Scale bars = 100 µm). (B) Quantification of Ki67 signal intensity normalized to DAPI signal intensity in the DG of P5 WT and *Aag*$^{-/-}$ mice (n = 3), data represented as mean ± SEM; statistical analysis performed using one-tailed Student's t-test. (C) Signal intensity of 5-hydroxymethylcytosine (5hmC) per cell binned by the number of phosphorylated H2AX (γH2AX) foci in the CA3 regions of the P5 hippocampi from WT and *Aag*$^{-/-}$ mice (n = 3). Data represented as median and quartiles, statistics performed using two-tailed Student's t-test; ns – nonsignificant ($p>0.05$), *** $p < 0.001$.

**Figure S2. Temporal gene expression analysis in WT and *Aag*$^{-/-}$ prefrontal cortex.** (A-C) Volcano plots depicting deferentially expressed genes (DEGs) in *Aag*$^{-/-}$ compared to WT pre-frontal cortex (PFC) at: (A) postnatal day 5 (P5), (B) 6 weeks (6W) and (C) 6 months (6M). (D) Top five biological processes (BP) gene ontology (GO) terms determined by the Gene Set Enrichment Analysis (GSEA) for DEGs in *Aag*$^{-/-}$ compared to WT PFC. (E) Heat map of the expression levels of genes differentially expressed in the PFC of *Aag*$^{-/-}$ mice compared to WT at all tested ages (P5, 6W, and 6M). Color scale is representing change in the gene expression depicting log2 fold change relative to the mean. (F-G) RTqPCR analysis of: (F) Ublcp1 and (G) Aag mRNA levels in in the P5 and 6M WT and *Aag*$^{-/-}$ hippocampi (n = 4). Data represented as mean ± SEM, significance determined by two-way ANOVA with Šídák´s multiple comparison test; ns – nonsignificant ($p>0.05$), **** $p < 0.0001$.

**Figure S3. Analysis of *Gabra2*, *Pttg1* and *Ublcp1* expression in the hippocampi of WT, *Aag*$^{Tg}$, and *Aag*$^{-/-}$ mice.** (A-C) RTqPCR analysis of: (A) Gabra2, (B) Pttg1, and (C) Ublcp1 mRNA levels in the hippocampus of P5 WT, *Aag*$^{-/-}$, and *Aag*$^{Tg}$ mice (n = 4). Data represented as mean ± SEM, statistics performed using on-way ANOVA with Holm-Šídák´s multiple comparison test: ** $p < 0.01$; *** $p < 0.001$; **** $p < 0.0001$.

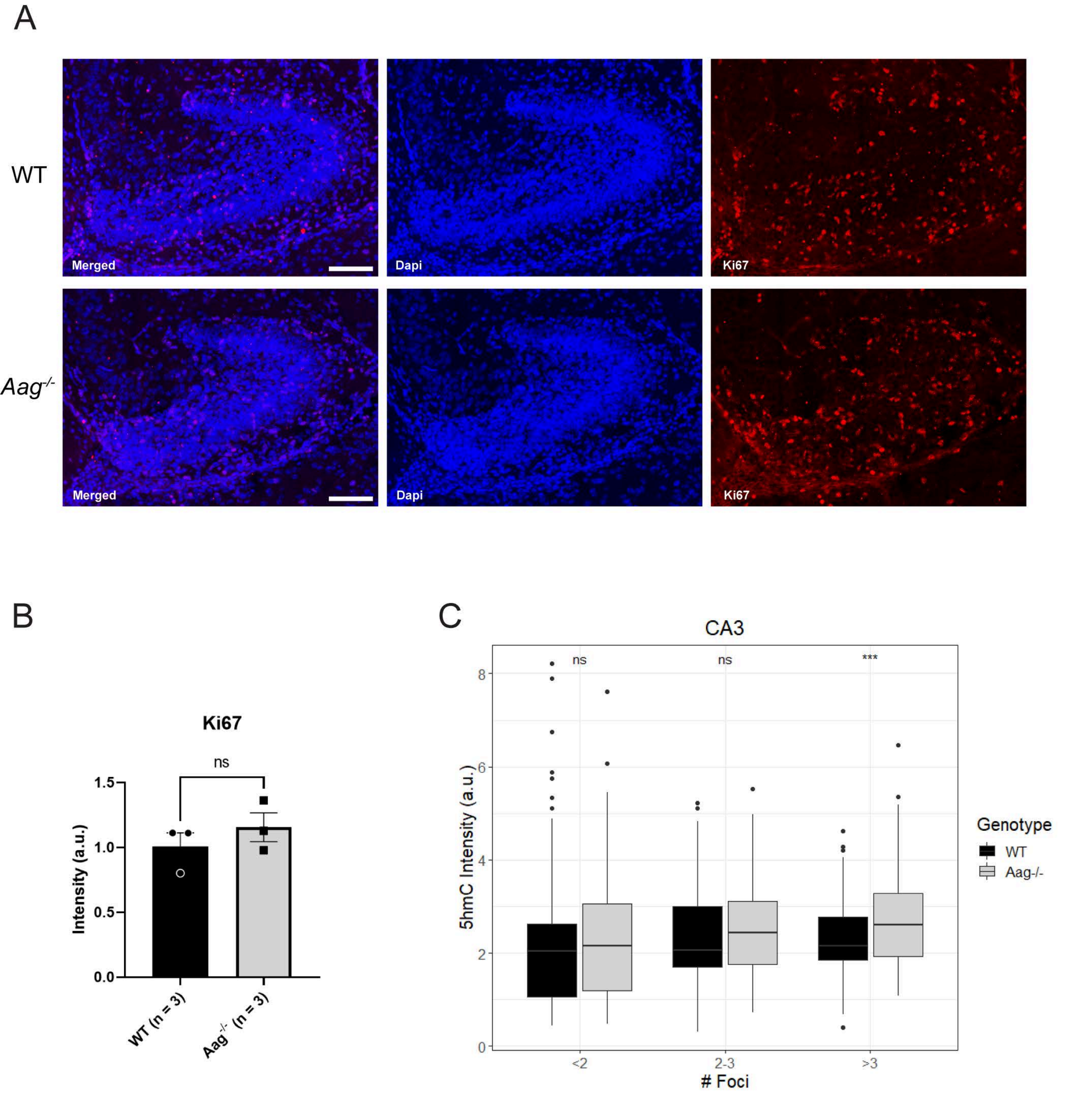
A
WT
Aag-/-
Merged
Dapi
Ki67
B
Ki67
ns
1.5
1.0
0.5
0.0
Intensity (a.u.)
WT (n = 3)
Aag-/- (n = 3)
C
CA3
ns
ns
***
8
6
4
2
0
5hmC Intensity (a.u.)
<2
2-3
>3
### Foci
Genotype
WT
Aag-/-

**Figure S1**

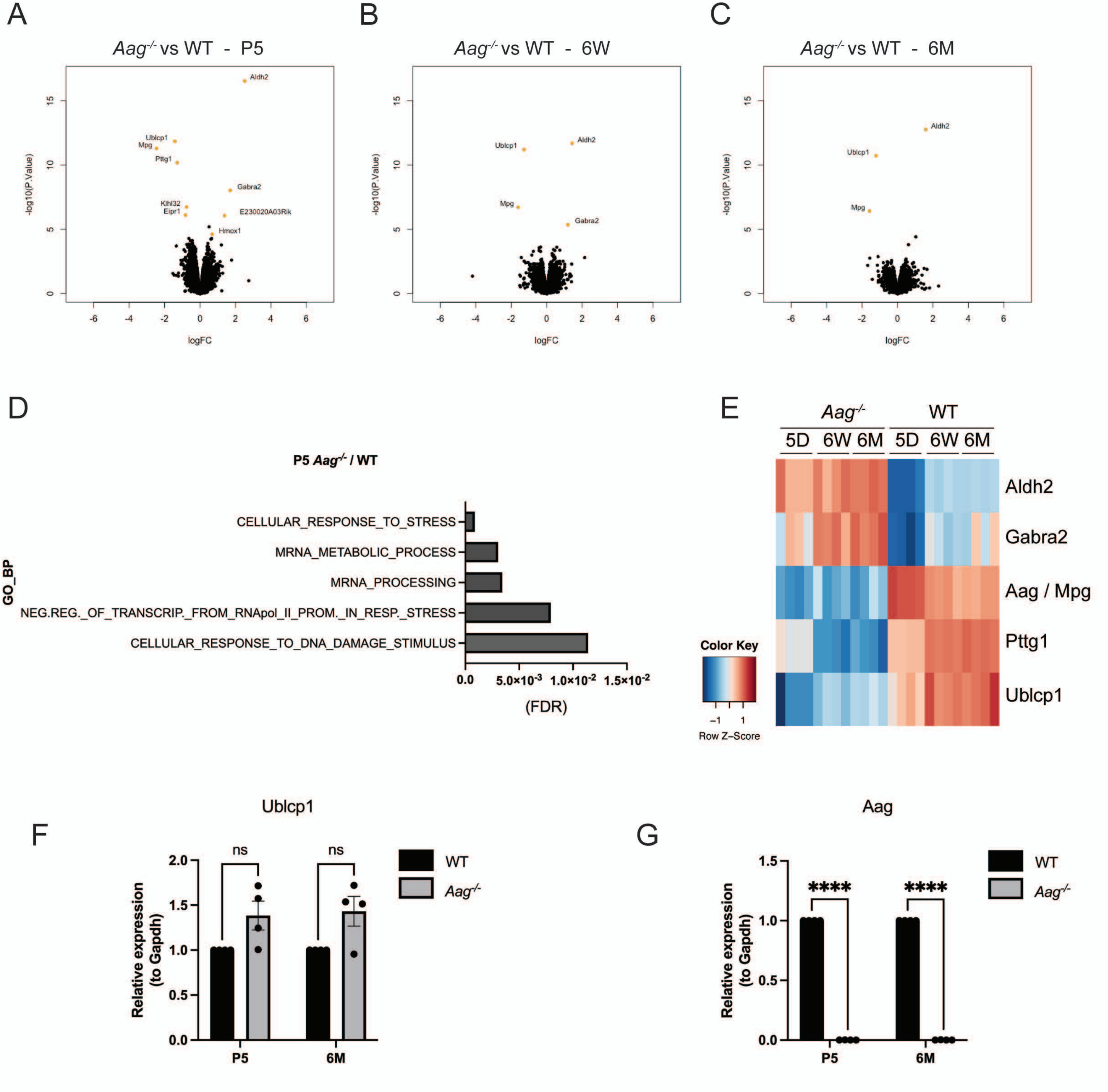
A
Aag-/- vs WT - P5
Aldh2
Ublcp1
Mpg
Pttg1
Gabra2
Klhl32
Eipr1
E230020A03Rik
Hmox1
-log10(P.Value)
logFC
B
Aag-/- vs WT - 6W
Ublcp1
Aldh2
Mpg
Gabra2
C
Aag-/- vs WT - 6M
Aldh2
Ublcp1
Mpg
D
P5 Aag-/- / WT
GO_BP
CELLULAR_RESPONSE_TO_STRESS
MRNA_METABOLIC_PROCESS
MRNA_PROCESSING
NEG.REG._OF_TRANSCRIP._FROM_RNApol_II_PROM._IN_RESP._STRESS
CELLULAR_RESPONSE_TO_DNA_DAMAGE_STIMULUS
0.0
5.0×10-3
1.0×10-2
1.5×10-2
(FDR)
E
Aag-/-
WT
5D 6W 6M 5D 6W 6M
Aldh2
Gabra2
Aag / Mpg
Pttg1
Ublcp1
Color Key
−1 1
Row Z−Score
Ublcp1
F
ns
ns
WT
Aag-/-
Relative expression (to Gapdh)
P5
6M
Aag
G
****
****
WT
Aag-/-
Relative expression (to Gapdh)
P5
6M

**Figure S2**

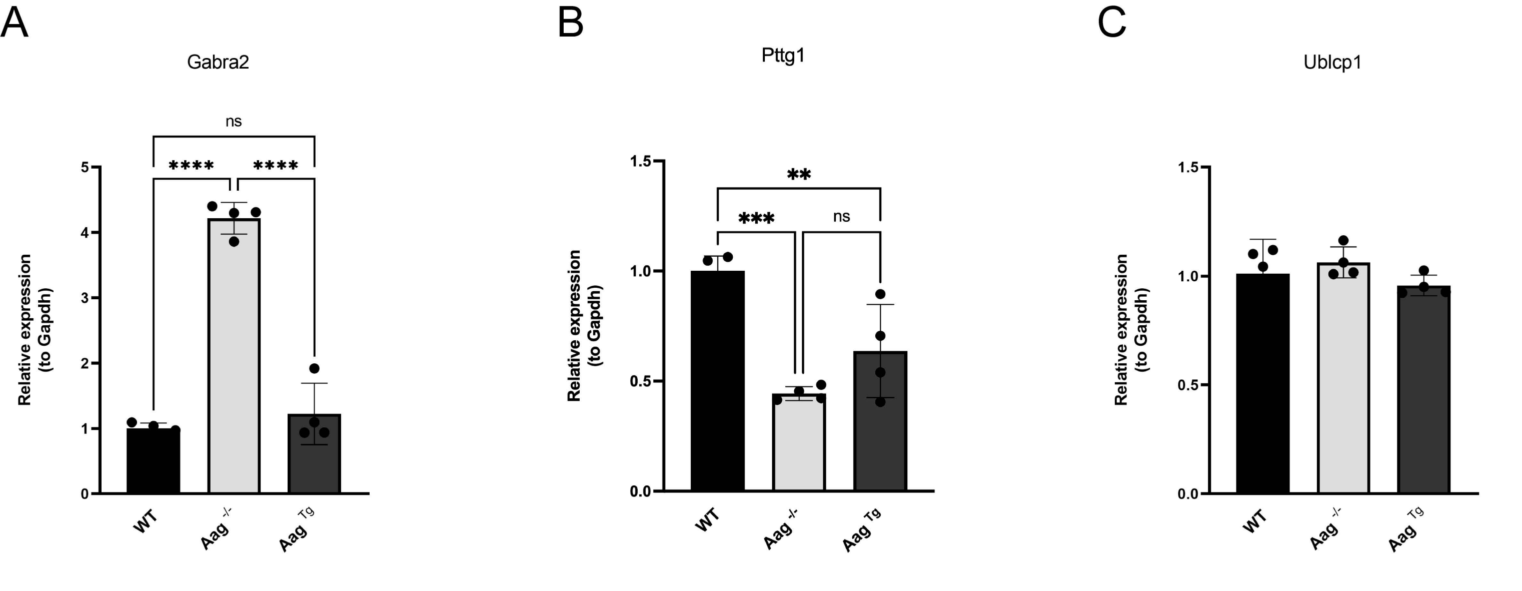
A
Gabra2
ns
****
****
Relative expression (to Gapdh)
0
1
2
3
4
5
WT
Aag -/-
Aag Tg
B
Pttg1
**
***
ns
Relative expression (to Gapdh)
0.0
0.5
1.0
1.5
WT
Aag -/-
Aag Tg
C
Ublcp1
Relative expression (to Gapdh)
0.0
0.5
1.0
1.5
WT
Aag -/-
Aag Tg

### Figure S3
